# Expanding the pulse-reserve paradigm to microorganisms on the basis of differential reserve management strategies

**DOI:** 10.1101/2022.02.24.481838

**Authors:** Ferran Garcia-Pichel, Osvaldo Sala

## Abstract

The pulse-reserve paradigm (PRP) is central in dryland ecology, although traits of microorganisms were not explicitly considered in its inception. We asked if the PRP could be reframed to encompass organisms both large and small. We used a synthetic review of recent advances in arid land microbial autoecology combined with a mathematically explicit theoretical model. Preserving the PRPs original core of adaptations by reserve building, the model considers differential organismal strategies to manage these reserves. It proposes a gradient of organisms according to their reserve strategies, from nimble responders (NIRs) to torpid responders (TORs). It predicts how organismal fitness depends on pulse regimes and reserve strategies thus explaining organismal diversification and distribution. After accounting for scaling phenomena and redefining the microscale meaning of aridity, it becomes patent that the PRP is applicable to microbes, and that this modified PRP represents an inclusive theoretical framework working across life-forms.

---

Drylands, which encompass deserts, shrublands, grasslands and savannas, account for 40% of the terrestrial surface (Prăvălie, 2016), 30% of global carbon fixation (Field *et al*., 1998) and explain most of the interannual variability of the global carbon cycle (Poulter *et al*., 2014). In addition, drylands are home of 30% of the human population and to some of most vulnerable groups of people, who are often immediately dependent on resources from the natural environment (Reynolds *et al*., 2007). However, theoretical frameworks of dryland functioning are scarce. This is particularly relevant when inclusive management of multiple ecosystem services is desired (Yahdjian *et al*., 2015, Hoover *et al*., 2020), including conservation of biological diversity as one such desired service. For example, to inform Managed Relocation efforts in the face of global climate change (Richardson *et al*., 2009), it would be relevant to have theoretical underpinnings of general applicability at the organism level. Additionally, microbial physiology, if not microbially-mediated processes, is underrepresented from the formulation of dryland ecological theory. A reasonable way to remedy this situation is to evaluate if current plant-based theoretical frameworks would accommodate the inclusion of microbes. The pulse-reserve paradigm (PRP) constitutes arguably the most important and enduring paradigm in dryland ecology since its postulation by Noy-Meir (Noy-Meir, 1973) almost 50 years ago. It holds that dryland ecosystems undergo cyclical dynamics in which rare rainfall events trigger a flow of resources, mostly carbohydrates and nutrients from reserves (seeds or perennial plant organs) to an existing plant. During the pulse, the plant grows new leaves and roots devoting most of the acquired resources through photosynthesis and absorption from the soil to new growth. Next, the plant starts sending resources to reserve organs that are replenished towards the end of this pulse to prepare for dormancy between pulses. As a new cycle starts with the next pulse, a new flow from reserves to growth will be triggered. The PRP was largely based on keen observations of the climatic rigors typical of drylands and on insightful naturalistic observations of plant life strategies and it has been subject to subsequent refinements and derivations relating to specific aspects like the importance of response thresholds (Schwinning *et al*., 2004), functional plant types (Ogle & Reynolds, 2004), and interactions with nutrient pulses (Collins *et al*., 2014). It remains one of the most cited works in the field, though microbes and microbial processes were under-represented at inception. It is “plant-centric” (Collins *et al*., 2014). And yet, the roles that microbes and their assemblages (microbiomes) play in dryland ecology have become ever more patent during the last decades (Bashan & de-Bashan, 2010). Not only are microbes influential as agents of disease, for which there is a long history of recognition, but they are now understood to be ubiquitous, sometimes intricately integrated partners in mutualisms with both plants and animals. They are also responsible for much of the biogeochemical cycling, and, in the face of limits to plant cover and productivity imposed by aridity, they can take on some basic ecosystem processes such as primary productivity and soil stabilization. Logically, efforts to bring microbial processes into the fold of the PRP eventually followed (Collins *et al*., 2008, Collins *et al*., 2014, Šťovíček *et al*., 2017). As so much in microbial ecology, however, these contributions were based on analyses of microbial communities and microbial processes. However, they did not directly consider differential adaptations in particular microbes, i.e, trait-based analyses (Martiny *et al*., 2015) in a way that parallels the core auto-ecological foundation of the Noy-Meir PRP for plants. The microbially based contributions thus remain a self-contained black box that offers none of the interpretive granularity that can be ascribed to plant communities. For example, the time scale for pulse responses is known to be much shorter for microbial than for plant processes (Collins *et al*., 2014), but while we can explain differential plant response times and thresholds on the basis of plant rooting depth, no such granularity can yet be achieved for microbes.

A successful strategy of scientific progress has been the borrowing of pieces of theory and their mathematical models from sister disciplines. Early on, concepts borrowed from the cost-income theory from economics were applied to the evolution of resource allocation between leaves and roots in plants of arid environments (Orians & Solbrig, 1977). Noy-Meir developed his PRP simultaneously under the intellectual influence of the International Biological Program (Boffey 1976). The cost-income theory is thus implicitly embedded in the PRP. Organisms need to utilize resources in the form of stored nutrients and energy to support a shift from quiescent to active state, to prepare for an eventual safe return to dormancy, and to mitigate environmental insults during the inactive state. Plants deploy new roots and leaves to capture resources after a rainfall pulse; their seeds germinate based on carbon reserves accumulated pre-emptively during the times of plenty. The PRP predicts that the resources acquired during the pulse will be enough to replenish storage to be able to restart the cycle at the onset of the next pulse, and result in some non-zero net growth during the pulse. Variability of the duration of the pulses determines whether enough resources are acquired to fulfill the needs. Short pulses may end up in the consumption of more resources than can be acquired, leading to organismal or population demise. Long pulses will certainly offset the initial investment and yield a positive effect on storage. Thresholds for shifts from quiescence to active state interacting with the length of pulses thus determine the outcomes for individuals and populations.

It is the intention of this contribution to probe if the core tenets in the PRP find applicability in microbial biology, a discipline that has become much more conspicuously important for the ecology of drylands than might have been surmised at the time, and one that now has experienced at least some advances to attempt such an exercise. Recent contributions show that it is indeed possible to attain general theoretical frameworks that are applicable to both micro-and macroorganisms when they are derived from basic, universal core tenets, as is the case with the principles of biological seed banks (Lennon *et al*., 2021). Broadening the PRP beyond plants has been very important in guiding integrative research in drylands and developing new tools for holistic dryland management. Our central questions are: 1) Is the PRP applicable to microbes at large? 2) Is there a universal PRP that can be applied from microbes to plants? and 3) What are the organismal adaptations in plants and microbes that allow them to manage different sized pulses? Since the PRP was first developed with plants in mind, we highlight plant-microbe contrasts in the process. Here, we introduce a new conceptualization: the importance of reserve management style as the basis for organismal adaptations to cope with pulsed resource availability. We frame it with a mathematical model, evaluate its predictions in the light of knowledge of organismal biology, and explore the consequences and applicability of the generalized paradigm.

## Modelling the PRP: nature and consequences of pulsed resources

We reframed the PRP to make it generically applicable but preserving its basic tenets. Succinctly, biological activity in drylands is strongly determined by pulses of water availability (Noy-Meir, 1973), and this challenge is met by organisms through the accumulation of reserves during the pulse that are then used to power transitions into and out of dormancy. The reframing is still based on the tenet that all organisms base their response to pulses on the accumulation of reserves, but we introduce the concept that organisms can differ in their reserve management style along a continuum between two strategies. At one end of the continuum, what we call “Nimble Responders (NIRs)” are organisms that transition swiftly in and out of pulse utilization mode. To achieve this, however, they allocate a certain proportion of their metabolism and resources to maintain a constant, constitutive physiological readiness for inter-pulse conditions, and to ensure that metabolic systems are inherently hardy and protected at all times. Such level of constant readiness comes at the cost of building permanent, constitutive reserves, thus depressing their growth potential (and rate) from a theoretical maximum; they are inherently slow growers. At the opposite end, “Torpid Responders (TORs)”, while they do allocate portions of their resources as reserves to prepare for quiescence, they only do so as a pulse nears an end. These can be understood as late-pulse investments. Consequently, they can allocate recently acquired resources fully to growth processes during much of the pulse unconstrained by allocation to reserves, and they grow swiftly during it. Because of their low level of constitutive readiness, the lag times for transition to dormancy and back into growth among TORs are long, in that reserves need to be mostly synthesized from scratch towards the end of a pulse and then retrofitted into growth-promoting biomass at the beginning of the next pulse. To continue the financial metaphor, NIRs can be understood as conservative investors, focused on certainty of returns that are not high but are frequent, while TOR would be akin to risky investors whose strategy relies on high returns that occur rarely. It is important to realize that specific organisms will fall within the continuum between the extremes exemplified in the NIR and TOR acronyms. The distinction will be most properly used when comparing the relative position of an organism along this continuum, and probably most useful when comparing the relative character of pairs of organisms.

An explicit mathematical model based on the ideas in the previous paragraph is presented in Supplementary Materials, the main formulations of which are gathered in BOX 1. Its input parameters, outputs and predictions are discussed in the following paragraph.

The duration of the pulse is divided into three periods: transition from quiescence, growth phase and transition to quiescence. The level of “constant readiness” for quiescence is gauged by the parameter α, which constitutes the central parameter in the model, defined and calculated as the proportion between reserve biomass and growth-yielding biomass maintained during the growth phase in the pulse. It remains constant within it, and growth must be positive, the level of constant readiness will vary between 0 minimally, and maximally approach 1. Organisms approaching the NIR end with high constant readiness will have high α, and TOR-like organisms will be characterized by low α. The level of investments in reserves needed to ensure successful transitions in and out of dormancy is gauged by the parameter, β, which is defined and calculated as the ratio of growth-yielding biomass to reserve biomass reached at the end of the pulse. Organisms that do not reach β by the end of a pulse will not be viable at the next iteration. Nothing prevents an organism to invest very heavily in β, and its maximal values are principally not constrained, but if these reserves are retrofitted into growth structures at the next pulse with reasonable efficiency (i.e., above 30%), β will typically take values below 3. The instantaneous growth rate (µ) that is possible at any one time during the growth phase in a pulse is dependent on α, being depressed from its maximal value by the resources allocated to constant readiness. The net yield of growth, defined as the ratio of growth enabling biomass at the end of a pulse to that at its beginning is, however, dependent on both α and β. The dynamics of growth-enabling biomass and reserve biomass during a pulse under this framework model are depicted in Fig. 1A. A comparative depiction of the dynamics of growth during a pulse for end-member organismal types is in Fig. 1B. Again, NIRs are characterized by low µ, but high α, while TORs by high µ but low α.

**Fig. 1.**
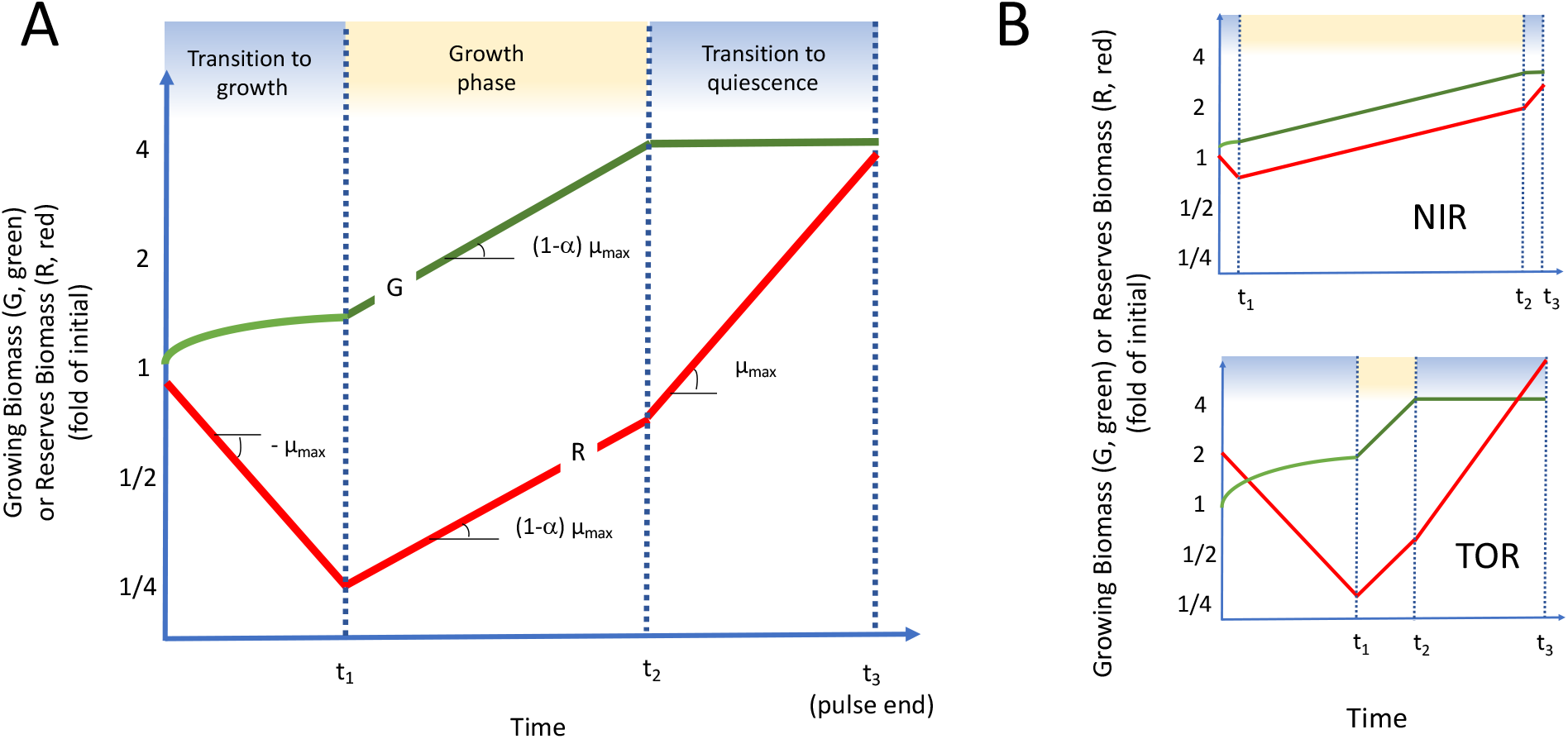
A: Modelled dynamics of growth-yielding biomass (G) and reserves (R) during a pulse of resources with explicit transitions between quiescence and growth and back to quiescence. Important parameters are intrinsic maximal growth rate (µ_max_), the ratio of biomass allocated constantly to reserves over that allocated to growth during growth (constant readiness, α), and the ratio of biomass allocated to support transitions over that allocated to growth at the end of the pulse. Times for transition are dependent on the previous parameters. B: Differential strategies of management of reserve investment result in distinct functional adaptations: NIRs and TORs. NIR organisms invest conservatively in preparation for quiescence (high α), and can respond swiftly (short transition times), at the expense of lower growth rate (μ), whereas TORs invest with higher risk (low α), need longer transition times but can grow fast during favorable times. Explicit mathematical model equations are in Supplementary Materials.

Relevant model outputs are the duration of the transition periods and growth phase, the net new growth yield after completion of a pulse cycle, and the minimal duration of a pulse required for viability. Their formulation as a function of input parameters is in BOX 1. The duration of the growth phase is determined by the duration of the pulse minus the duration of the necessary transitions. Pulse duration must exceed the duration of transitions for organisms (or populations) to attain net growth. While the point has been made that transitions are energetically costly for both plants (Gremer & Sala, 2013) and microbes (Schimel, 2018), they are also costly in terms of time, particularly when active time is at a premium. Hence, we also define a *minimal pulse duration* that allows for completion of the necessary transitions. Should a pulse last less than this minimal duration, viability will be compromised and the resuscitation strategy will fail.

The model predicts that: 1) Transition times decrease with increasing constant readiness reserves (α) and increase with increasing end-of-pulse reserves (β); only in the case in which α tends to β, they tend to 0, which biologically corresponds to the case in which virtually all reserves needed for transition are always present. 2) The minimal duration of the pulse needed for viability decreases with increasing constitutive readiness (Fig. 2). 3) Increasing the amount of end-of-pulse reserves (*β*) lengthens minimal pulse duration for viability.

**Fig.2.**
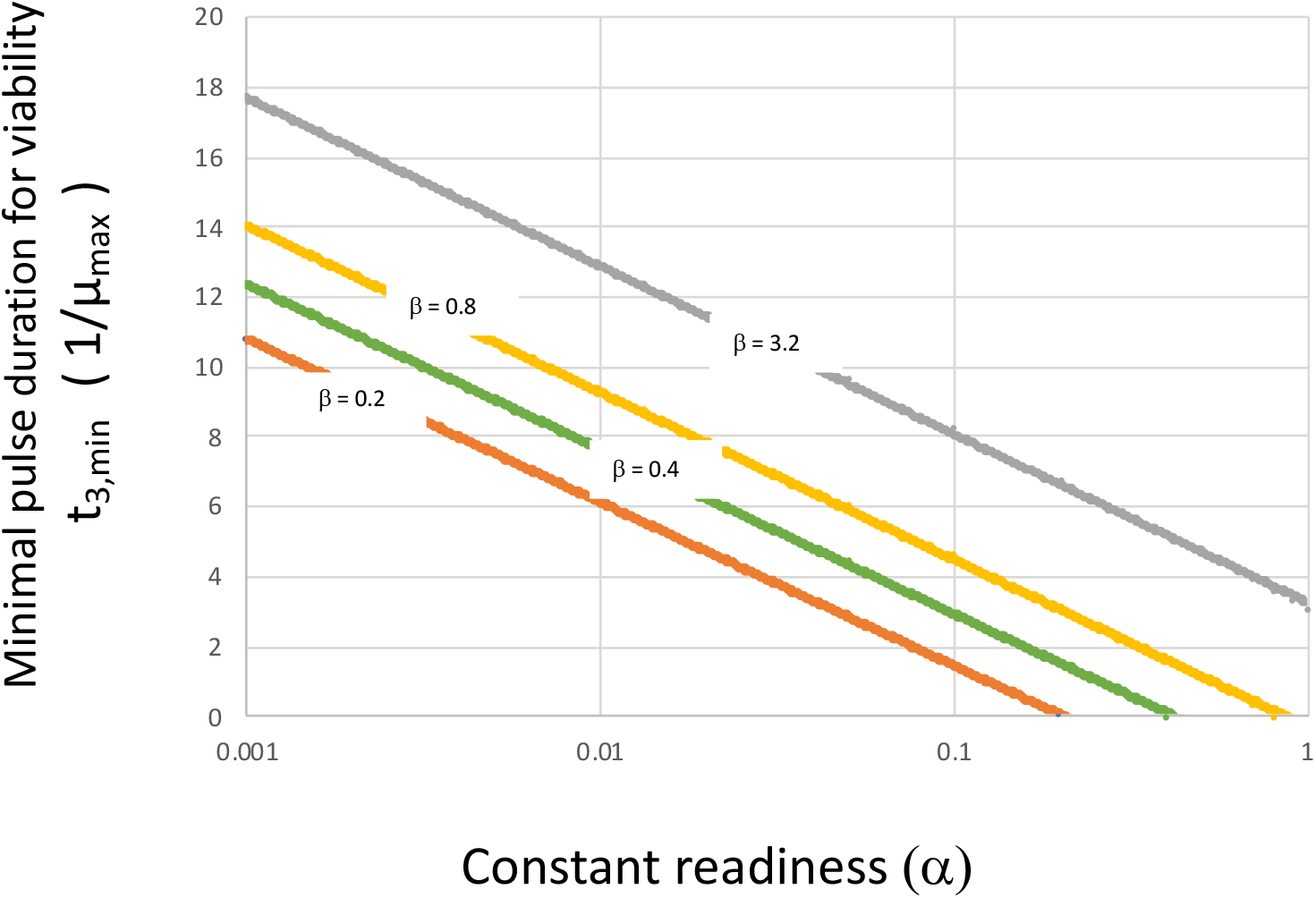
Minimal pulse duration for viability (t_3, min_, expressed as multiples of the inverse of µ_max_) as a function of level of constant reserve investment (constant readiness, α) for various values of late-pulse reserve investment (β).

In terms of growth yields as a function of pulse duration (Fig. 3), the model predicts that: 1) The more an organism tends to the NIR end-member, the more it will benefit from a regime of short pulses whereas tending to the TOR end, will benefit from longer pulses. 2) Investments in end-of-pulse reserves (β) result in advantages in growth yield regardless of pulse duration, as long as the minimal pulse duration is exceeded. 3) The return of late-pulse investment becomes increasingly negligible for very long pulses. In general terms, the constant readiness of NIRs is only beneficial when pulses are short, while late-pulse investments matter more when pulses are intermediate in duration. When pulses are so long as to lose their “pulsed” character neither type of adaptation is beneficial, consistent with the PRP tenet that the gathering of reserves is a characteristic selected for only under pulsed regimes of growth.

**Fig. 3.**
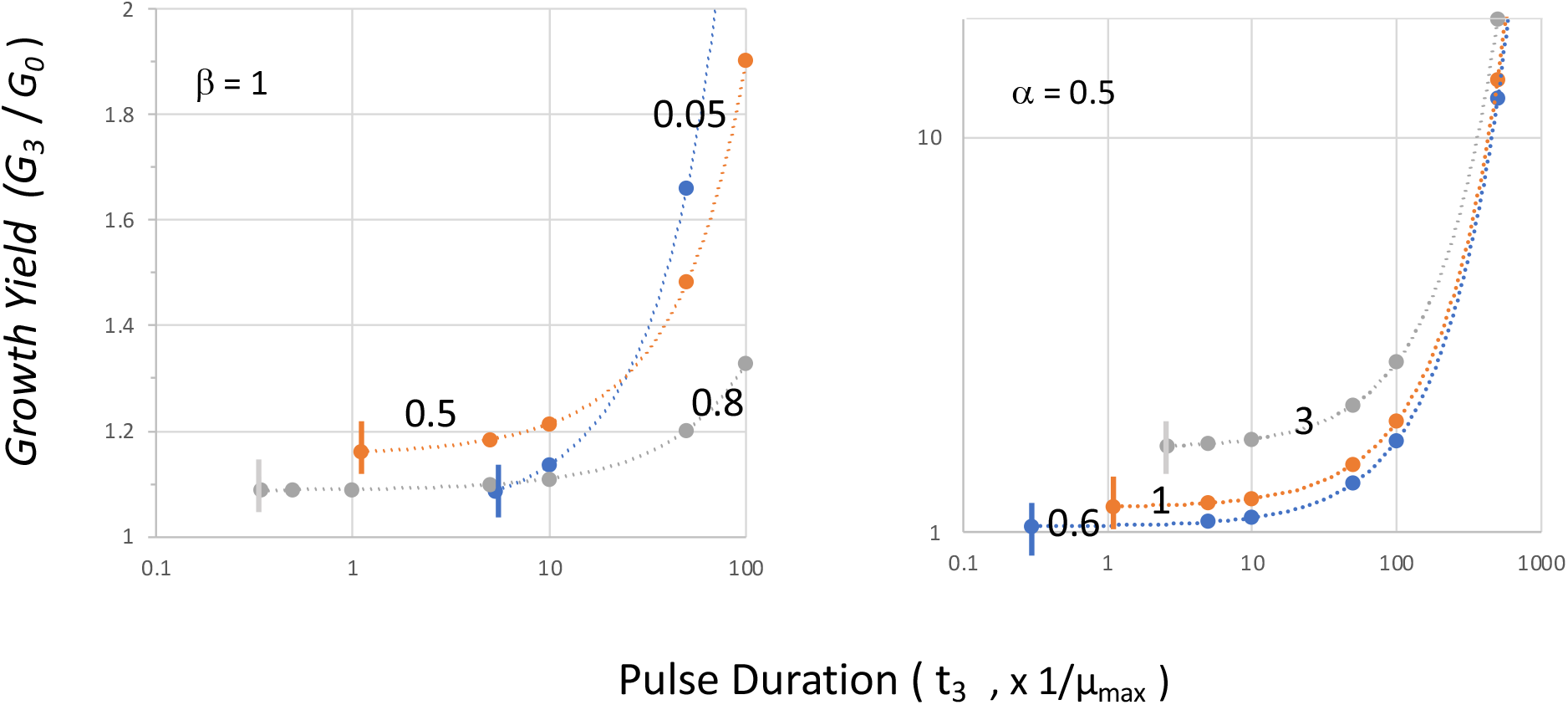
Dependence of growth yield (defined as fold increase in G over the duration of a pulse) on pulse duration (expressed as multiples of the inverse of µ_max_), for various values of constant readiness α (left) and late-pulse investments β (right). Vertical bars indicate minimal pulse duration as per Fig. 2. Only moderate yields are ever attained with high α, but positive growth is attained even with very short pulses. Higher yields can be attained by increasing β, but at the expense of higher thresholds for viability, and the beneficial effects become irrelevant for very long pulses.

While we have expressed pulse duration, rather unintuitively, as the inverse of µ_max_, these units have a biological meaning. In exponential growth 1/µ_max_ represents the time it takes for a population to grow by a factor of *e*. It is related to the (minimal) doubling time, *D*, as *D* = ln2/µ_max_. So, in biological terms, NIR organisms will have an advantage in pulses that are shorter than some 6 *D* (below some 10 times 1/ µ_max_; Fig. 2). For bacteria, while there is considerable variability, average doubling times are around 5 h (Weissman *et al*., 2021). Thus, for bacteria, recurrent pulse durations shorter than a day will benefit only NIRs, and above that, TORs will do better, but “pulses” exceeding a week, will not benefit either type. For plants, the analogous whole-plant relative growth rates (RGR) vary between 0.1 and 0.4 d^-1^ (Price & Munns, 2020), and we can take 3 days as a typical doubling time. NIR plants will benefit from (soil) water pulses shorter than 18 days, but that will be insufficient for TOR plants. Pulses of water availability that exceed many months will not yield differential benefits to either type.

Importantly, however, all these predictions are derived from very simple, generic tenets that do not involve specific plant functional traits, so that they are therefore universally applicable. Pulse duration/size bestows benefits differentially between NIRs and TORs, and its variability can principally drive organismal specialization, in turn supporting differentiated ecosystem-level outcomes, which is a fundamental characteristic of the PRP (Schwinning *et al*., 2004). The predictions thus logically imply that shifts in pulse size distribution range will impact the functional types of organisms that can effectively grow, the spatial extent of their populations and their dynamic fate under climates changing in time or space. Experimental evidence consistent with this prediction has been presented: imposed treatments with decreased average rain pulse size result in swift and very marked shifts in community composition of primary producers of biological soil crusts, favoring cyanobacteria over mosses (Reed *et al*., 2012), and favoring some species of cyanobacteria over others (Fernandes et al. submitted). Similarly, the predicted shift in plant species composition under regimes of differing pulse size have been presented (Jones *et al*., 2016). Traits that might implicate differential management of reserves among losers and winners, however, were not directly quantified in any of those cases.

On the other hand, it is not difficult to find principally NIR-like and TOR-like cases among arid-land plants. For example, perennial grasses such as *Bouteloua gracilis* that dominate enormous areas of North America have shown a remarkably fast response to precipitation pulses. Not only leaf water status and photosynthesis but also root growth (Lauenroth *et al*., 1987) rapidly shift in a few hours after a rain event into an active state, returning to quiescence a few days after. These grasses invest constitutively in non-structural carbohydrate reserves (Menke & Trlica, 1983), lignocellulose, and in hardy crown tissue structures containing buds and tillers that together allow fast onset of growth. Yet, they produce only moderate amounts of seeds. Annual grasses that dominate California grasslands, by contrast, are examples of TOR-like plants: comparatively, they grow swiftly without conspicuous investments in structures dedicated to improving survival under inter-period conditions, but they do so only during rather large pulses of precipitation (Huenneke & Mooney, 2012). Towards the end of the pulse their physiology quickly turns into the all-out production of reserves in the form of large amounts of seeds. The ultimate question for us remains: is there evidence that microorganisms also fit in this framework? Since its premises are only a pulsed nature of growth, the reliance on reserves as strategies, and the differential management of reserves by specific types, the question translates essentially to finding evidence that microbes rely centrally on reserve-building and to find a functional differentiation between NIRs and TORs among them.

## Microbial traits in drylands

Before we embark on the attempt to characterize traits of dryland microbes, we should consider that climatic definitions were not made with them in mind. Some microbes greatly exceed the perceived limits of temperature and water availability that underlay climatic cutoffs (Rothschild & Mancinelli, 2001, Schulze-Makuch *et al*., 2018), and spatial environmental variation within a single climatic region can create non-compliant microniches. Texturally coarse soils of mesic climates, for example, unable to retain meteoric water inputs for very long do often sustain microbial communities typical of deserts in temperate climates (Smith *et al*., 2004). In hyper-arid settings, the presence of surface pebbles that locally retard soil water evaporation allow the formation of hypolithic microbial communities, embedded in an otherwise barren landscape (Schlesinger *et al*., 2003, Warren-Rhodes *et al*., 2006). Thus, paradigms that may ensue from the study of microbial adaptations to aridity will find full applicability outside of climatically understood drylands and will fail to apply in some instances within them. One should apply the concept of aridity in agreement with the implied outcome of biological activity hindrance by water scarcity. Still, several representative characteristics of dryland plant communities do in fact find parallels in their microbial co-inhabitants. A direct evaluation of differential genomic capabilities as well as comparative surveys of microbial distributions reveal that microbial communities and their member microbes are uniquely adapted to arid conditions, as are desert plants and animals. Global distributional surveys of soil bacteria clearly point to a compositional differentiation in drylands from those in other climates (Angel *et al*., 2013, Neilson *et al*., 2017, Delgado-Baquerizo *et al*., 2018, Schulze-Makuch *et al*., 2018), a pattern that seems to involve preferentially rare species, i.e. species with a narrow-niche (Bickel & Or, 2021). These are traits they share with plant and animal communities. In fact, for a few specific microorganisms, we have sufficient evidence that their distribution is constrained to arid environments, implying a maladaptation to other climates (Hector & Laniado-Laborin, 2005). And yet, as it is true for microbes in most environments, we know relatively little about the specific nature of these differential adaptations, given that most microbial ecology efforts are not geared to establishing the “natural history” of particular microbes, their auto-ecology, but rather to establish emergent properties of microbial communities taken as black boxes. But it is precisely these organismal auto ecological traits that comprise the central pillars of the PRP. At most, changes in community composition can be used to infer trait-based patterns assuming that some traits have a phylogenetic correlation. But this exercise is fraught with perils, in that phylogeny seems to underpin certain traits more strongly than others and also differentially so among major microbial groups (Martiny *et al*., 2015). The opposite concept is that of convergent evolution in which similar traits evolved along different evolutionary paths both in plants (Orians & Solbrig, 1977) and microbes (Garcia-Pichel & Wojciechowski, 2009)

Microbes in arid conditions do lead a life characterized by rare pulses of activity embedded in a regime of suspended animation (Lennon & Jones, 2011, Schimel, 2018). Because of the short generation times of microbes, community composition can change dramatically even within the duration of a single pulse, based on the resuscitation of dormant, rare species (Aanderud *et al*., 2015). This implies that selective forces acting on traits that sustain transitions from dormant to active states are at play. Metagenomic (Le *et al*., 2016) and metatranscriptomic (Rajeev *et al*., 2013) analyses of arid soil bacterial populations speak of the crucial importance of the capacity to repair genetic damage accumulated during inactive periods for an effective recovery from dormancy. Activation of DNA repair seems to be the absolute priority inasmuch as they constitute the very first wave of gene expression upon microbial rehydration (Setlow, 2007). The studies carried out on two desiccation-resistant model bacteria, *Deinococcus* (Cox *et al*., 2010) and *Chroococcidiopsis* (Billi *et al*., 2000) tell us a similar story about the relevance of damage repair. Adaptations to avoid photodamage, such as the synthesis of passive sunscreen compounds (Garcia-Pichel *et al*., 1992), or the increase in cellular ploidy to attain genomic redundancy (Cox *et al*., 2010, Sukenik *et al*., 2012), also speak of the importance of adaptations to increase overall fitness by decreasing mortality during quiescence. While synthesis of these components is accomplished during active periods of growth, their contribution to fitness takes place under metabolic dormancy, when active repair does not take place. The sunscreen scytonemin, for example, is common in cyanobacteria that thrive under pulsed growth conditions, including those from soil crusts and epilithic environments (Garcia-Pichel & Castenholz, 1991), but has been recurrently lost through the evolution of the Phylum Cyanobacteria in clades that adapted to more constant environments (Garcia-Pichel *et al*., 2019). The decoupling of light-harvesting systems from reaction centers in desert microbial phototrophs constitutes another example of a preventative strategy (Bar-Eyal *et al*., 2015), and so does the migration towards environmental refugia in anticipation of desiccation (Pringault & Garcia-Pichel, 2004). Similarly, significant investment into spore formation (Setlow, 2007) is common in saprophytic fungi and in at least two phyla of bacteria, the Firmicutes and the Actinobacteria (Lennon *et al*., 2021). These examples clearly speak for an anticipatory, integrative coordination between the “system states” of quiescence and active growth, a shift that can occur rather rapidly at the physiological and genetic levels (Rajeev *et al*., 2013, Oren *et al*., 2019). Thus, microbes adapt to the pulsed nature of their environment, rather than to either the active or inactive phases in it. This brief review of microbial life strategies suggests that a preemptive accumulation of reserves (in the form of cellular machinery and energy reserves to fuel repair upon activation, or as investment in costly secondary metabolites to avoid damage) could well play a role in the swift and effective use of short pulses of hydration and activity; and may in fact underpin the adaptation of microbes to life in arid lands.

### Size differentially affects how microbes and plants cope with a pulsed environment

The small size of microbes imposes constraints and opportunities that differ significantly from those affecting plants. Relevant processes scale with size (L), oftentimes because of the differential scaling of body volume (L^3^) to surface (L^2^). In any attempts to test how the PRP applies to microbes, one should take those effects into account. For example, because the usefulness of suncreens depends directly on the thickness of the teguments they are laid onto, plants can use suncreens much more efficienty for photoprotecion than microbes can (Gao & Garcia-Pichel, 2011). Because conductive heat transfer scales inversely with size (Planinšič & Vollmer, 2008), very small organisms equilibrate temperature with its surrounding virtually instantaneously. Temperature homeostasis simply cannot be a microbe’s forte: microbes must evolve to endure temperature shifts unscathed instead. Mass transport is very strongly scale sensitive. At small scales, a microbe’s environment becomes essentially stagnant, diffusion becoming the main mass transport vehicle; but diffusion’s effectiveness scales with L^-2^. This is why bacteria do not need internal transport structures, and why metabolite exchange among microbes at short range is quite effective, requiring little metabolic effort. Desiccation can be understood in similar physical terms, as the diffusional loss of water in a gradient of water potential, and hence fractional water loss scales with L^-2^ (Jakubczyk *et al*., 2012, Carrier *et al*., 2016). Again here, microbial cells will tend to equilibrate with their immediate surroundings virtually instantaneously and microbial cells will be as wet as their environment is. Attempting to keep water inside a microorganism by retarding evaporation using non-diffusive teguments (as some plants do using cuticular waxes) is principally a lost cause.

An important caveat for the generalizations above, however, is that aggregation of microbial cells into larger, coherent structures such as colonies or biofilms can reverse these physically imposed opportunities and limitations. Nutrient supply can be limited by diffusion when microbes come together into mm-sized aggregates or biofilms (Garcia-Pichel *et al*., 1999). When they do, they typically create microenvironments around such aggregates (Garcia-Pichel & Belnap, 2001) because diffusion is insufficient to release their metabolic byproducts. In some extreme cases, such macroscopic aggregates can cause turbulent transport of nutrients through motility (Fenchel & Glud, 1998), increase the albedo and equilibrium temperature of soil surfaces (Couradeau *et al*., 2016), or retain moisture for some time after a rain event (Scherer & Zhong, 1991). Macroscopic aggregates of microbes must be independently evaluated for the relevance of such effects.

### Do microbes also use activity pulses to gather reserves in preparation for quiescence?

Microbes do produce organic and inorganic storage reserves, typically in the form of polymers that help them avoid high cellular turgor pressure: polyglucose (glycogen, starch) and poly-β-hydroxyalkanoates (PHAs) are carbon reserves. These type of reserves are prevalent among microbes (Mason-Jones *et al*., 2021). More specialized polymers, like cyanophycin, store carbon and nitrogen, and polyphosphates store inorganic phosphorous (Kolodny *et al*., 2006). Colorless sulfur bacteria can even accumulate electron donors like polymeric sulfur and acceptors like nitrate (Schulz *et al*., 1999). Polymerization and mobilization of all of these reserves is responsive to pulses in availability of the respective nutrient (Dawes & Senior, 1973), in patterns suggestive of a prioritization of storage over growth when resource depletion is imminent (Mason-Jones *et al*., 2021). The gathering of reserve polymers is generally considered a major physiological trait enabling microbial dormancy and the formation of seed banks (Lennon & Jones, 2011). A genetic ability to produce abundant reserves, correlates in specific microbes with resistance to stress conditions (Madueño *et al*., 2018) and can demonstrably support resuscitation from dormancy (Klotz *et al*., 2016). Recent studies of gene expression in field populations of the cyanobacterium *Microcoleus vaginatus* from soil crusts during wetting/desiccation pulses showed the importance of reserve compounds in the context of overall gene regulatory acclimation: genes for cyanophycin and polyphosphate synthesis were expressed heavily during the “growth”, wet phase (see Fig.1), whereas those for the mobilization of both glycogen and polyphosphate reserves were strongly upregulated with the onset of desiccation, in the “transition to quiescence phase” of Fig.1. Direct measurements and metabolic simulation have shown that *M. vaginatus* allocates more fixed carbon to polymers constitutively than other cyanobacteria (Jose *et al*., 2018), indicating that this microbe tends to be a NIR type, which is consistent with field experimentation (Fernandes et al, submitted). Both these facts are consistent with the pulse-reserve hypothesis as it pertains to carbon and nutrients, the reserves accumulated during wet times allowing the preparation for successful dormancy (i.e., anticipatory regulation) during drought. Other microbes are known for the accumulation of specific resources in preparation for desiccation or exit from quiescence phases (Klotz & Forchhammer, 2017). The soil actinobacterium *Rhodococcus* upregulates the formation of ectoin (a compatible solute), and proteins to detoxify oxygen stress (LeBlanc *et al*., 2008). Such investments can be significant. UV sunscreen compounds, like cyanobacterial scytonemin, can make up investments in the order of several percentage points of a cell’s dry mass, becoming useful in comparison to active repair mechanisms only during quiescence (Gao & Garcia-Pichel, 2011) and its biosynthesis is enhanced by desiccation (Fleming & Castenholz, 2007). Glycogen or PHA can be accumulated to more than half of a cell’s dry weight (Klotz & Forchhammer, 2017, Madueño *et al*., 2018). Such level of accumulation would speak for associated values of the parameter β in our model easily exceeding unity.

The developmentally sophisticated formation of desiccation-induced spores (Sukenik *et al*., 2015) (Setlow, 2007), a strategy shared by fungi and several bacterial phyla, represents perhaps an extreme case of all-out late-pulse allocation to reserves (typical of TORs with low *α*) that enables extremely long quiescence periods. The trade-off here is that a relatively long new pulse with copious resources is needed to allow for an equally complex process of germination (Stewart *et al*., 1981, Dworkin & Shah, 2010) before new resources can be effectively tapped, as predicted by our model. Thus, as in plants, not only do microbes prepare for dormancy through the gathering of resources, but examples of microorganisms adapted to either short or long pulses that manage these reserves differently can also be found.

### Soil segregates between microbial NIRs and TORs

In Noy-Meier’s framework, soil acts as a water repository that significantly extends water availability beyond the precipitation pulse, for organisms that can tap it, (Fig. 4) and is a central consideration in the original and subsequent refinements of the paradigm (Ogle & Reynolds, 2004). As one proceeds down the soil profile, water content suffers from diminished inputs (infiltration inputs will require larger pulses) but benefits from more moderate evaporative losses, so less water reaches deep layers but there it lingers over longer periods. So, pulse duration increases with soil depth (Sala *et al*., 1992). This differential distribution of water in the soil profile translates into a force for plant diversification, where NIR-tending functional types specialize in tapping surficial resources whereas TOR-like shrubs utilize mostly deeper water (Jackson, 1999). But, how does this translate to soil microbial populations? We have seen how in principle the duration of a pulse for a microbe will tend to be as long as the water is available in its immediate surroundings. Undoubtedly, system responses that rely on microbially mediated processes are much faster than those mediated by plants (Austin *et al*., 2004). Consistent with long known empirical evidence (Linn & Doran, 1984), soil microbial activity should follow the dynamics of soil water content along the profile and in time (Fig. 4). We can thus expect that microbes will function for short periods at the surface (NIR territory), and for longer periods as they move down the profile (increasingly TOR turf). In fact, it is those spore formers in the Firmicutes and the Actinobacteria (TOR-like organisms) that tend to dominate zones deep in the soil profile (Fierer *et al*., 2003), where high water content episodes are rare but once they happen, they last for long time. In soil surface communities such as biocrusts, by contrast, TOR spore-formers are typically exceedingly rare under normal conditions (Garcia-Pichel *et al*., 2003), although veritable blooms of spore-formers can be brought about by long flooding of these crusts (Karaoz *et al*., 2018). Typical soil-surface communities, like biocrusts, are inhabited by communities that respond swiftly, within minutes, to pulses (Scherer & Zhong, 1991, Garcia-Pichel & Belnap, 2001). The point has been made that the ecosystem contributions to carbon cycling of such communities is particularly important under regimes of short pulses (Cable & Huxman, 2004). Further, even within surface communities, plant-interspaces are prone to experience more intensely pulsed conditions than the soil under plant canopies. Trait-based comparisons of microbial communities in these two contrasting environments from an arid setting show a significant enrichment of spore-forming microbes in soil under plants (more TOR-like in our framework), even when generic traits to resistance to desiccation were more enriched in the interspace soils (Goberna *et al*., 2014).

**Fig. 4.**
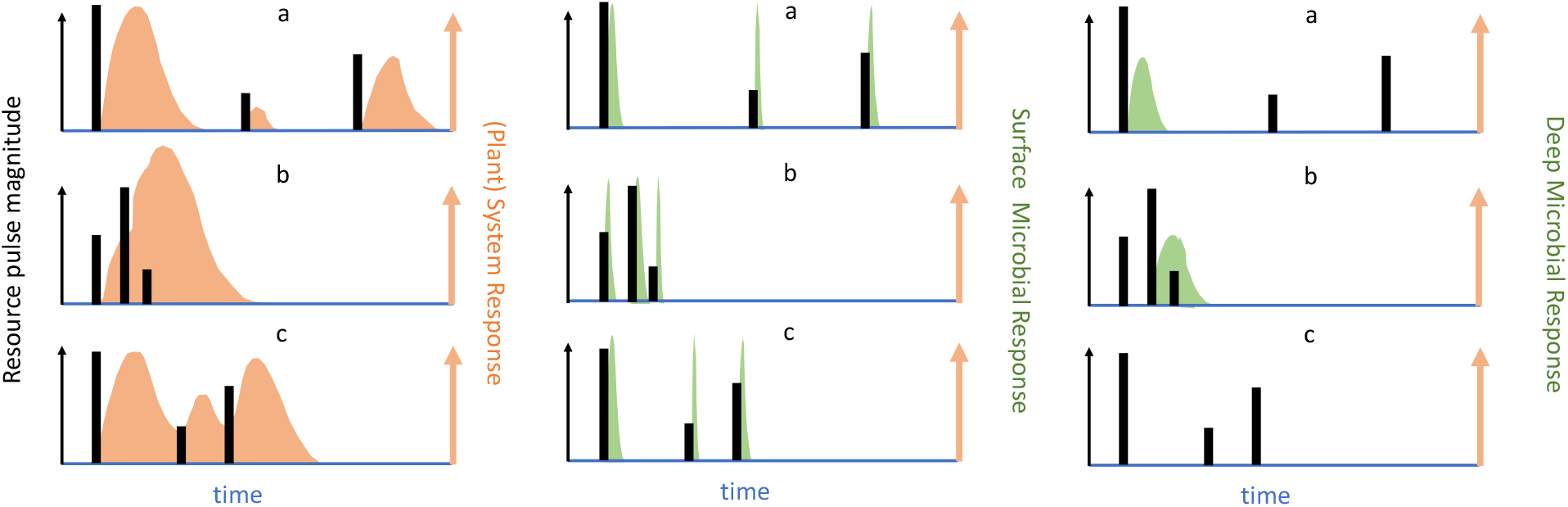
Noy-Meier’s concept of the plant-system responses (shaded areas) to pulsed water resources (black bars) in the left panel, with its translation to microbial responses for NIR-like microoorganisms close to the soil surface (middle panel) and for TOR-like microorganisms deeper in the soil profile (right panel). Panels a to c represent different pulse regimes.

The mechanisms by which this is accomplished differ between plants and microbes. The shorter generation times of microbes allow this to take place at the level of community assembly, rather than just as evolutionary eco-physiological diversification. In fact. experiments show that even short-term variations in aridity can exert rather swift microbial community shifts in a variety of settings (Rothrock Jr & Garcia-Pichel, 2005, Castro *et al*., 2010, Barnard *et al*., 2013, Liu *et al*., 2019). In a mesoscale experimental set up, soil crust cyanobacterial communities suffered stark shifts in organismal dominance within a few years of either a decrease in overall precipitation or a 2 month delay in the onset of the rainy season (Fernandes *et al*., 2018), promoting dominance of NIR-like, drought-resistant *M. vaginatus* over other cyanobacteria. These dynamic compositional shifts find consistent parallels in the geographical distribution of the same cyanobacterial taxa along varying climates (Fernandes *et al*., 2021). The patterns reviewed above indicate that while the effects on diversification operate through alpha-diversity in plant communities, they do so through beta-diversity along the soil profile in microbial communities. In sum, both in plants and in microorganism, the pulsed nature of water distribution in the soil is a force for niche partitioning and evolutionary diversification, as postulated by Noy-Meir.

### Formally testing the model

We have sought here to assess the NIR/TOR model for consistency with current organismal knowledge. While the phenomenology reviewed is consistent with its predictions, it is only indirectly and qualitatively so, and it encompasses only the few organismal types whose biology was informative enough. It will be important in the future to test the model more formally, quantitatively, and generally. A conceptually simple experimental test could make use of imposed variations in pulse duration and pulse number as the cumulative pulse time is kept constant, as has been previously used to assess effect at the community or ecosystem level (Jones *et al*., 2016)(Fernandes et al, submitted), but with a focus on quantitative determination of the main parameters in the model: µ, α and β. The expectation here is that experimental increase in pulse frequency and reduction in pulse duration will result in changes in microbial community composition that favor the presence of organisms with elevated α and lower µ. Alternatively, one could seek a correlative validation by comparing the average NIR vs. TOR character in locales selected along geographical gradients of pulse-size distribution. Naturally the duration of experimental treatments will have to allow for differences in generation times between plant and microbes. In the case of microbes, determination of α, β and µ can be best performed using relevant isolates in culture. For plants, single-species specimens for analyses can be easily obtained from the field (Gremer & Sala, 2013). Quantification of α and β is rather straightforward using biochemical analyses for known reserve compounds (Del Don *et al*., 1994, Martínez-Vilalta *et al*., 2016) during the appropriate time (growth phase for α, or initial quiescence phase for β).

Organismal traits can also be principally derived from (meta)genomic information directly from natural microbial communities. For example, genomically-coded information can be used to indirectly gauge habitat breadth (Barberán *et al*., 2014), maximal growth rates (Weissman *et al*., 2021), or nutritional modes (Chen *et al*., 2021) of specific phylotypes. Further, comparative analyses of genomic complexity in the genes underlying particular traits provides proxies for the relative importance of a given trait in a given setting (Cao *et al*., 2020). In some cases, organismal traits can be assigned to particular phylotypes on the basis of sequence similarity to known organisms (Goberna *et al*., 2014, Couradeau *et al*., 2019), making it principally possible to derive organismal traits from the most common form of community data analyses (i.e, 16S rRNA amplicon sequencing), although this requires assumptions of phylogenetic conservation that are far from guaranteed (Martiny *et al*., 2015). Metagenomic approaches are much preferred in this context. To test predictions regarding the NIR-TOR continuum, genomic/transcriptomic proxies for its three main parameters (µ, α and β) can be readily envisioned. Ribosomal codon use bias can be used to gauge the maximal growth rates commonly realized by an organism in its habitat, as it reflects adaptations of the protein synthesis machinery to those maximal rates. This will gauge our parameter µ (rather than our µ_max_, which defines a theoretical maximal growth rate for any organism under a given set of conditions and resources). While the relative importance of reserves in a given microbe can potentially be ascertained by the presence, relative abundance and complexity of the genes that code for their synthesis and mobilization (Rajeev *et al*., 2013), the direct derivation of proxies for α and β, is probably more complex, if not impossible, since the genetic underpinning of constitutive (α) and end-of-cycle (β) reserves is likely shared. An assessment of relative gene expression levels during growth and transition phases will be required, which will necessitate the use of time-resolved (meta)transcriptomics during actual pulse experiments.

## Conclusions and Outlook

Yes, the PRP works from microbes to plants. However, the mechanisms that evolved to cope with an environment characterized by resource pulses followed by prolonged periods of quiescence are different for microbes and plants. Size determines the mechanisms that make organisms thrive in this highly variable and extreme conditions. We do find examples approaching NIR and TOR organisms in both plants and microbes. In many regards, Noy-Meir’s vision of the pulse-reserve dynamics shaping the natural history of desert organisms is applicable to microbes, which show a diversity of adaptations much richer than he could have envisioned when he lumped microorganisms under the term “poikilohydric”. These adaptations are in many regards convergent with those of plants, although they are often modulated by scaling constraints. The PRP clearly holds across levels of biological organization for carbon reserves and for specialized structures when understood as a function of strategies to manage necessary reserves (Fig 5). The consequences of a pulsed water regime for functional diversification hold across organismal boundaries but operate largely through organismal diversity in plants and more intensely through community assembly in microbiomes.

**Fig. 5.**
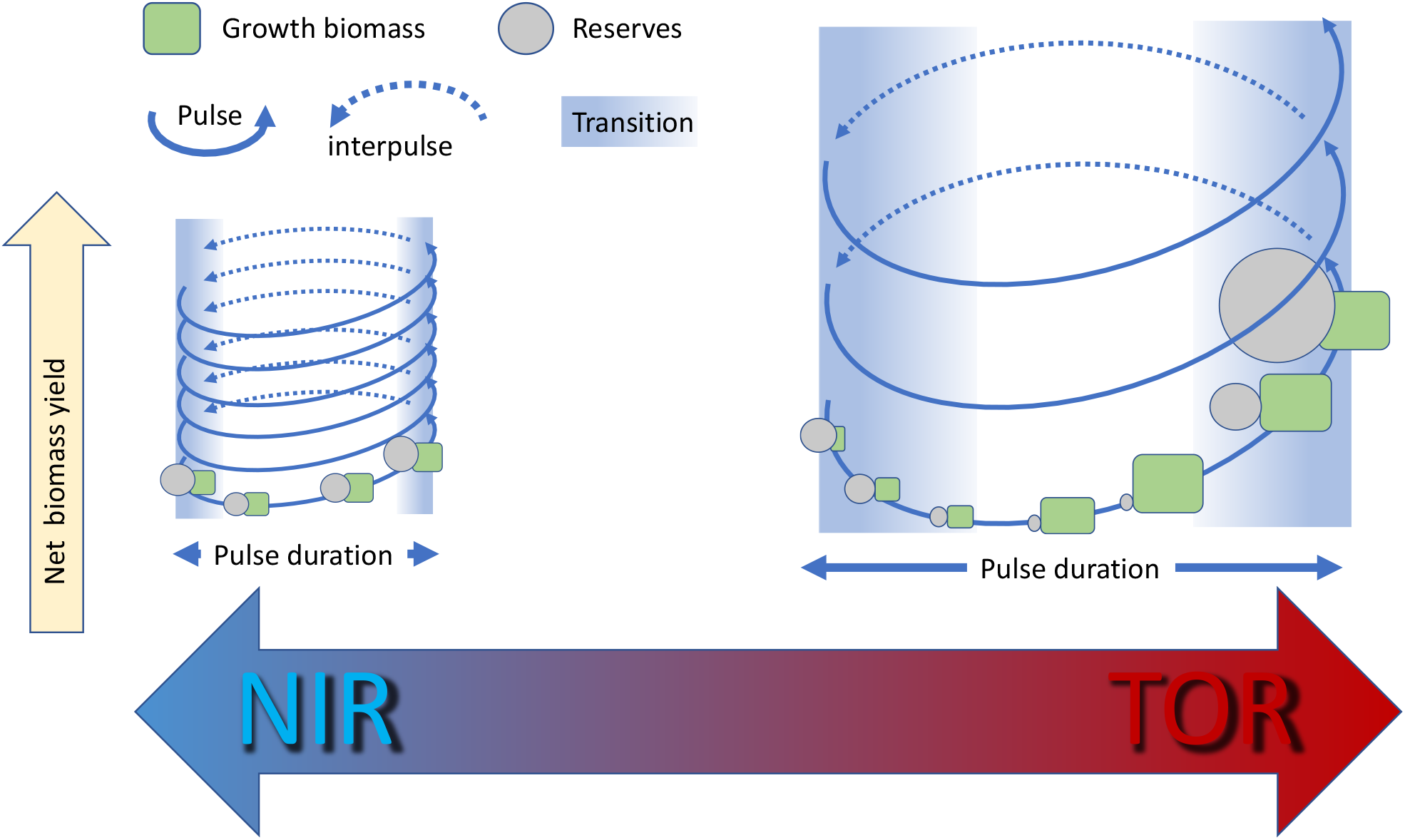
Conceptual abstraction of the organismal strategy continuum for the management of reserves that result in optimal returns under pulsed activity regimes of varying pulse length. Most organisms will find themselves in the continuum between nimble responders (NIR) and torpid responders (TOR).

While the current formulation of our model suffices to predict many of the predicted outcomes of the PRP, it does not explicitly include two aspects of arid land organismal adaptations that may *a priori* be considered relevant: the capacity for acquisition of water reserves and the length of periods in between pulses. Acquisition of water reserves serves the purpose of effectively lengthening the pulse duration and is a common adaptation among dryland plants like cacti and euphorbias. The issue with water reserves is much tougher for microbes than for plants, because of the size constraints discussed above, although extracellular polysaccharide investments may slow rates of water loss (Roberson & Firestone, 1992, Chenu, 1993, Hart *et al*., 1999). This is clearly so in microbes that build macroscopic thalli like *Nostoc commune* (Scherer & Zhong, 1991), or in microbial assemblages that attain sufficient mass, but are unlikely to make much of a difference for single, typical microbial cells. This may very well be behind the well-known fact (Potts, 2001, Alpert, 2005) that only small organisms (or small independent parts of plants, like seeds) can truly be anhydrobiotic, that is survive a temporary loss of all (or most) of their water, since reliance on anhydrobiosis is the only viable solution for small beings. Drought resistance in macro-organisms, by contrast, is always dependent on preventing water loss. Increasing length of pulse inter-periods will decrease the capacity of dormant cells to resuscitate as they accumulate environmental insults that will have to be repaired. In our framework, this will mean that organisms faced with long droughts will experience a decrease in the efficiency with which reserves can be retrofitted into growing biomass, which we had assumed to remain constant and high. This is likely to be countered by increases in *β* to make up for the loss in efficiency, which in turn will increase the minimal pulse size for viability, and thus likely affect NIR types more negatively than TOR types. These aspects should be given more detailed attention in future work.

Given that continental microbial ecosystems similar to those now present in drylands are known to have existed (Simpson *et al*., 2013, Beraldi-Campesi *et al*., 2014) and to have driven global biogeochemical cycles (Thomazo *et al*., 2018) since the mid-Precambrian, long before the advent of land plants, perhaps the fact that microbes follow the PRP should not come as a big surprise, since opportunity makes the thief. An apparently universal differential aspect of the biology of living organisms under arid regimes is that fitness under pulsed conditions in a background of water scarcity becomes a strong function of strategies to prevent death during periods of quiescence, whereas mechanisms geared towards promoting fast growth become less relevant than they might be in mesic environments. In other words, desert organisms, including microbes, can be expected to be hard to grow but hard to kill. Microbes probably found that out first, the hard way.

## Acknowledgements

The authors acknowledge the support of the National Science Foundation (NSF) through grants DEB 2025166 (Jornada Range LTER), DEB 2129537, DEB 1754106, New Phytologist Foundation and the ASU Global Drylands Center.

## BOX 1.

Formulations for growth dynamics and main output parameters

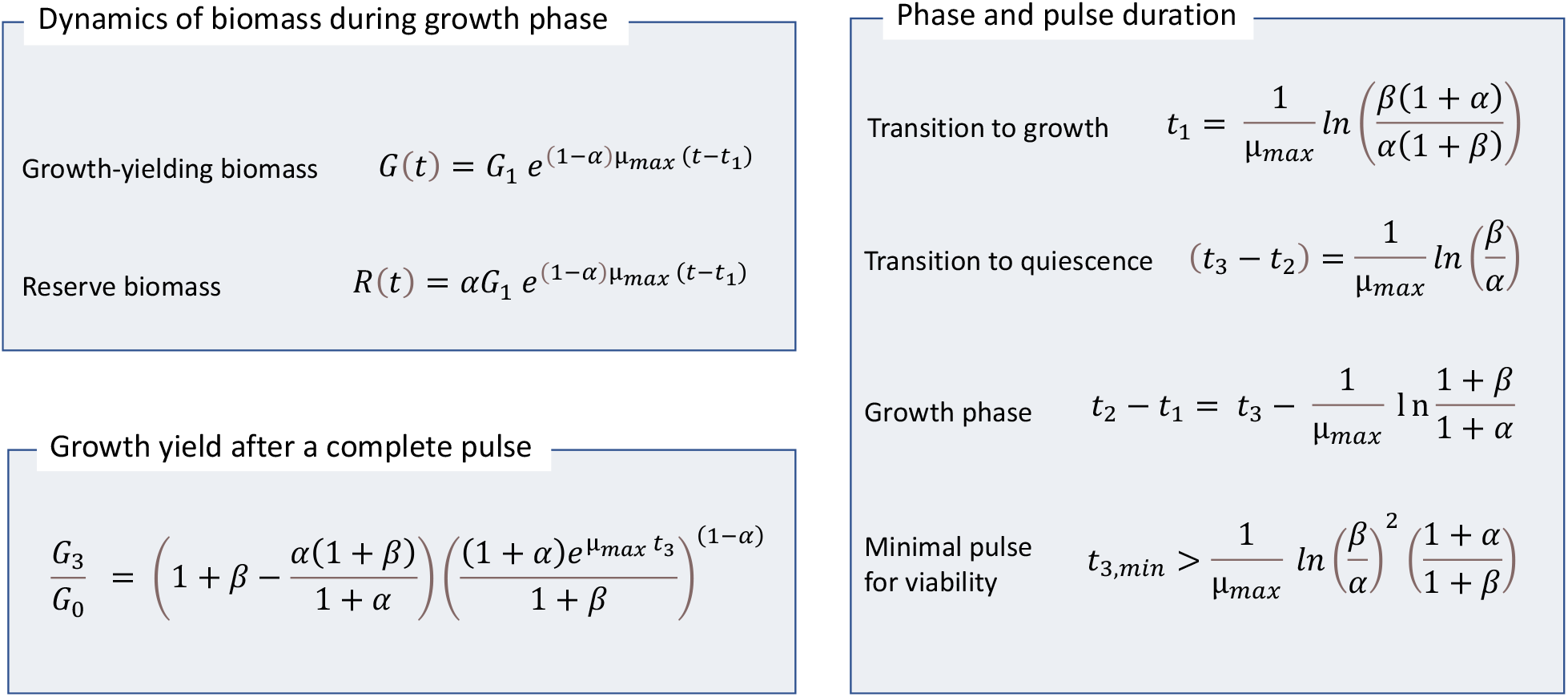

## SUPPLEMENTARY MATERIAL

### Explicit modeling of pulsed growth

#### 1. Growth phase

Let’s assume that during growth episodes a proportion of the total resources are converted into biomass capable of growth while the rest is allocated to reserves. Let us name the growing biomass *G* and the reserve biomass *R*. Let us also assume that growth is exponential. If no reserves are allocated at all, then growth is maximal, µ_max_, as determined by both inherent physiological constraints, and the growth dynamics will follow the differential equation:

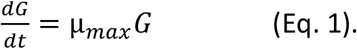

But if the proportion of biomass allocated to reserves is α, then G will instead follow

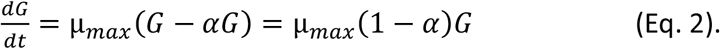

Thus, *G* will growth with a smaller, but also exponential, rate and

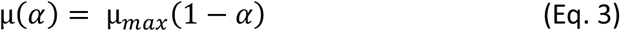

Note that because μ(α) > 0, *α* < 1

In integrated form Eq. 2 becomes:

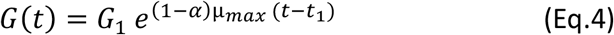

Where *t* is time, t_1_ is the start time of the growth phase, and *G*_1_ is the value of *G* at t_1_.

Since during the growth phase a constant proportion of G is allocated constitutively to resources, R = αG, in this phase

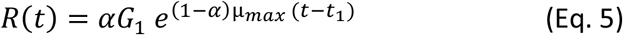

Hence the reserves also grow exponentially keeping pace with the growing biomass.

#### 2. Transition from quiescence into growth

Transitioning successfully from quiescent status to growth status requires resources that are proportional to the existing biomass *G*_*0*_ at the beginning of the transition. Let β be this proportion, so that *R*_0_= β *G*_*0*_. R is used to power this transition, including activities of damage repair, repurposing of resistance structures into growth-enabling structures and so on^1^. We can assume that the rate of use of R is maximal and equivalent to the inherent maximal growth rate µ_max_, R(t) in this phase thus decaying exponentially. The decay in R during this transition will be given by:

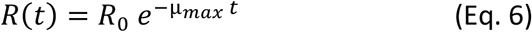

And it will proceed until R reaches the proportion of reserves typical of the growth phase, α. We note that by necessity α < β, 0 < α < 1, and β > 0.

When this is reached, at t_1_, the organism is ready to switch into growth mode, starting with a G_1_ that has increased from G_0_ by the new growth-enabling biomass additions from mobilized reserves. Thus:

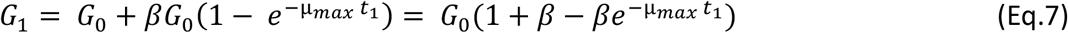

The end of the transition phase and the beginning of the growth phase occurs when *R*(*t*) = *α G*_1_. Substituting Eqs. 6 and 7 and solving for t, we find that

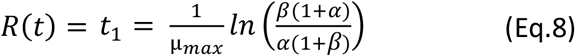

Thus, the duration of the transition depends inversely on the maximal growth rate, and on the proportions of biomass allocated to reserves constitutively, α, and at the beginning of the pulse, β. The transition time decreases with increasing α and with decreasing β. As β tends to α , t_1_ tends to 0.

#### 3. Transition from growth into quiescence

Upon sensing that the pulse is coming to an end, organisms must switch from growth mode into full reserve-gathering mode, so that by the end of the pulse, the proportion of reserves has increased from the constitutive α to the amount needed to support an eventual transition back to growth during a future pulse, β. This is accomplished by redirecting resources fully from growth to reserves, and growth stalls. After this time of reckoning, t_2_, G will remain constant at G_2_, and reserves will grow at full capacity following

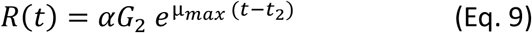

until they reach the necessary level, *βG*_*3*_(*=βG*_*20*_)

The anticipatory interval, t_3_-t_2_, needed to attain this must satisfy

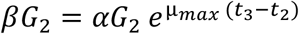

so that

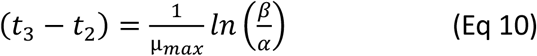

Thus, the duration of this anticipatory period, increases logarithmically with decreasing α and with increasing β. As α tends to β, it approaches 0.

#### 4. Integrated dynamics through the pulse

The total growth during a pulse of duration *t*_*3*_ comes from initial transfers from reserves and from actual new additions during the growth phase. The former is given in Eq. 7 and the latter can be calculated from the duration of this growth phase, [*t*_*2*_ - *t*_*1*_]. Eq. 8 gives us *t*_*1*_, and Eq 10 gives us *t*_*3*_*-t*_*2,*_ *so that*

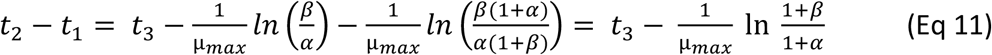

As α tends to β, [*t*_*2*_ - *t*_*1*_] approaches *t*_3_. As α decreases, [*t*_*2*_ - *t*_*1*_] decreases logarithmically with it.

It is then possible to express explicitly the growth during a pulse as a function of its duration, t_3_, given α, β, and μ_max,_

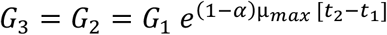

where *G*_*1*_ is given by Eq. 7, so that

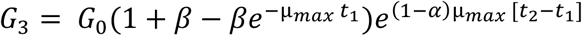

And where t_1_ and [t_2_-t_1_] are given in Eqs. 8 and 11, respectively, so that

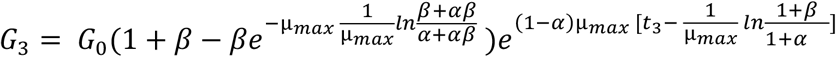

Making 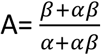 and 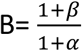, we get:

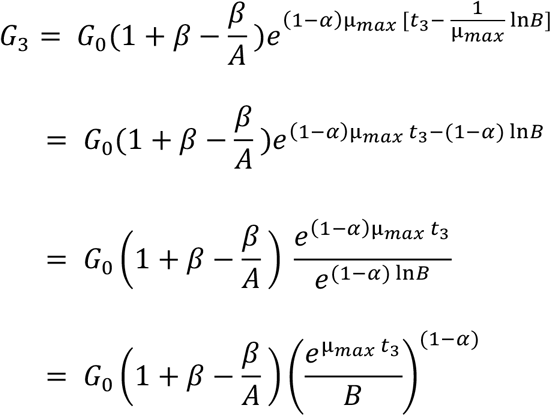

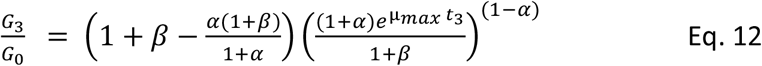

#### 5. Evaluation of minimal transition times

However, t_3_ must minimally allow for both transition periods to take place fully, or otherwise the proportion of reserves will have decreased below the level needed to allow a successful resuscitation at the next pulse. Thus, this minimum duration, t_*3,min*_, must be equal or exceed the sum of t_1_ and t_3_-t_2_, which are given by Eqs. 8 and 10, respectively, so that:

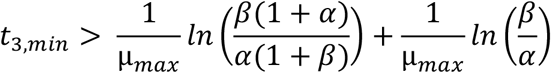

or

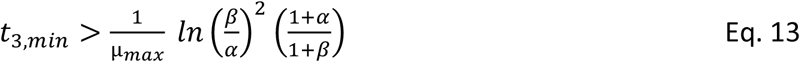

This can be conveniently evaluated with *t*_*3,min*_ expressed as multiples of 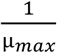 (see Fig. 2 in main manuscript. We see that the minimal duration of the pulse increases quasi-logarithmically with decreasing *α*, and that increasing *β* increases duration quasi-linearly. Also, that the minimal duration of the pulse becomes 0 only when *α = β*, and that keeping α high for any given *β*, minimizes transition times.

#### 6. Evaluation of net growth yield as a function of pulse duration

Equation 12 gives us the biomass attained during a pulse integrating the processes that occurred during growth and transition phases, *G*_*3*_, as a ratio to the initial biomass available *G*_*0*_ and as a function the pulse’s duration, *t*_*3*_, for a set of values of parameters *α*, *β*, and *μ*_*max*_. Again here, it is convenient to evaluate the dependencies expressing time as multiples of 1/*μ*_*max*_, which is related to the inherent maximal doubling time of the population, *D*, as *D*= ln(2)/*μ*_*max.*_

A graphical evaluation of the effect of low, medium and high values of α, the portion of the biomass dedicated to reserves constitutively, with fixed β, and considering the minimal transition times derived in the previous section can be found in Fig. 3 of the main manuscript.

For low values of α, copious growth is attained only with long pulses whose duration exceed tens of times 1/ μ_max_, and the minimal transition period for success is above five times 1/ μ_max._ Conversely a high α results in moderate but positive growth for short pulses, even with pulses as short as 0.3/ μ_max,_ but with low yields even for long pulses. Organisms with high α will outcompete those with low α if the pulses tend to be short, and vice-versa. This constitutes a biological adaptation strategy. Organisms that opt for low α can be called TOR and those that opt for high α , NIR.

One can also evaluate the influence of β, the portion of the biomass present as reserves at the beginning (and end) of the pulse. Fig. 3 in the main manuscript shows graphically the effects of varying β with fixed α. For very long pulses, beyond 100/ μ_max_ this has very little effect on yield. However, increasing β above unity increases the minimal length of the pulse to values well above 1/ μ_max_. Conversely, low β decreases this minimal threshold below 1/ μ_max_at the expense of significantly lowered yield. Thus, organisms with low β will do better in short pulses. Within the range of medium sized pulses, around 10/ μ_max_, large β results in optimal yields. With very long pulses strategies of varying β become rather irrelevant.

## Footnotes

1 We will assume that all reserves are being used to build growth-yielding structures, and that this happens with an efficiency of 100%

## Literature Cited

Aanderud ZT, Jones SE, Fierer N & Lennon JT (2015) Resuscitation of the rare biosphere contributes to pulses of ecosystem activity. Frontiers in Microbiology 6: 24.

Alpert P (2005) The limits and frontiers of desiccation-tolerant life. Integrative and Comparative Biology 45: 685–695.

Angel R, Pasternak Z, Soares MIM, Conrad R & Gillor O (2013) Active and total prokaryotic communities in dryland soils. FEMS microbiology ecology 86: 130–138.

Austin AT, Yahdjian L, Stark JM, Belnap J, Porporato A, Norton U, Ravetta DA & Schaeffer SM (2004) Water pulses and biogeochemical cycles in arid and semiarid ecosystems. Oecologia 141: 221–235.

Bar-Eyal L, Eisenberg I, Faust A, Raanan H, Nevo R, Rappaport F, Krieger-Liszkay A, Sétif P, Thurotte A & Reich Z (2015) An easily reversible structural change underlies mechanisms enabling desert crust cyanobacteria to survive desiccation. Biochimica et Biophysica Acta (BBA)-Bioenergetics 1847: 1267–1273.

Barberán A, Ramirez KS, Leff JW, Bradford MA, Wall DH & Fierer N (2014) Why are some microbes more ubiquitous than others? Predicting the habitat breadth of soil bacteria. Ecology Letters 17: 794–802.

Barnard RL, Osborne CA & Firestone MK (2013) Responses of soil bacterial and fungal communities to extreme desiccation and rewetting. The ISME journal 7: 2229.

Bashan Y & de-Bashan LE (2010) Microbial populations of arid lands and their potential for restoration of deserts. Soil biology and agriculture in the tropics,p.^pp. 109–137. Springer.

Beraldi-Campesi H, Farmer JD & Garcia-Pichel F (2014) Modern terrestrial sedimentary biostructures and their fossil analogs in Mesoproterozoic subaerial deposits. Palaios 29: 45–54.

Bickel S & Or D (2021) The chosen few—variations in common and rare soil bacteria across biomes. The ISME Journal 1–11.

Billi D, Friedmann EI, Hofer KG, Caiola MG & Ocampo-Friedmann R (2000) Ionizing-radiation resistance in the desiccation-tolerant cyanobacterium Chroococcidiopsis. Applied and environmental microbiology 66: 1489–1492.

Cable JM & Huxman TE (2004) Precipitation pulse size effects on Sonoran Desert soil microbial crusts. Oecologia 141: 317–324.

Cao H, Shimura Y, Steffen MM, Yang Z, Lu J, Joel A, Jenkins L, Kawachi M, Yin Y & Garcia-Pichel F (2020) The trait repertoire enabling cyanobacteria to bloom assessed through comparative genomic complexity and metatranscriptomics. MBio 11: e01155–01120.

Carrier O, Shahidzadeh-Bonn N, Zargar R, Aytouna M, Habibi M, Eggers J & Bonn D (2016) Evaporation of water: evaporation rate and collective effects. Journal of Fluid Mechanics 798: 774–786.

Castro HF, Classen AT, Austin EE, Norby RJ & Schadt CW (2010) Soil microbial community responses to multiple experimental climate change drivers. Appl Environ Microbiol 76: 999–1007.

Chen Y, Neilson JW, Kushwaha P, Maier RM & Barberán A (2021) Life-history strategies of soil microbial communities in an arid ecosystem. The ISME Journal 15: 649–657.

Chenu C (1993) Clay—or sand—polysaccharide associations as models for the interface between micro-organisms and soil: water related properties and microstructure. Soil Structure/Soil Biota Interrelationships,p.^pp. 143–156. Elsevier.

Collins SL, Sinsabaugh RL, Crenshaw C, Green L, Porras-Alfaro A, Stursova M & Zeglin LH (2008) Pulse dynamics and microbial processes in aridland ecosystems. Journal of Ecology 96: 413–420.

Collins SL, Belnap J, Grimm N, Rudgers J, Dahm CN, D’odorico P, Litvak M, Natvig D, Peters DC & Pockman W (2014) A multiscale, hierarchical model of pulse dynamics in arid-land ecosystems. Annual Review of Ecology, Evolution, and Systematics 45: 397–419.

Couradeau E, Giraldo-Silva A, De Martini F & Garcia-Pichel F (2019) Spatial segregation of the biological soil crust microbiome around its foundational cyanobacterium, Microcoleus vaginatus, and the formation of a nitrogen-fixing cyanosphere. Microbiome 7: 1–12.

Couradeau E, Karaoz U, Lim HC, Da Rocha UN, Northen T, Brodie E & Garcia-Pichel F (2016) Bacteria increase arid-land soil surface temperature through the production of sunscreens. Nature communications 7: 10373.

Cox MM, Keck JL & Battista JR (2010) Rising from the Ashes: DNA Repair in Deinococcus radiodurans. PLoS genetics 6: e1000815–e1000815.

Dawes EA & Senior PJ (1973) The role and regulation of energy reserve polymers in micro-organisms. Advances in microbial physiology, Vol. 10 p.^pp. 135–266. Elsevier.

Del Don C, Hanselmann KW, Peduzzi R & Bachofen R (1994) Biomass composition and methods for the determination of metabolic reserve polymers in phototrophic sulfur bacteria. Aquatic sciences 56: 1–15.

Delgado-Baquerizo M, Oliverio AM, Brewer TE, Benavent-González A, Eldridge DJ, Bardgett RD, Maestre FT, Singh BK & Fierer N (2018) A global atlas of the dominant bacteria found in soil. Science 359: 320–325.

Dworkin J & Shah IM (2010) Exit from dormancy in microbial organisms. Nature reviews microbiology 8: 890–896.

Fenchel T & Glud RN (1998) Veil architecture in a sulphide-oxidizing bacterium enhances countercurrent flux. Nature 394: 367.

Fernandes V, Giraldo-Silva A, Roush D & Garcia-Pichel F (2021) Coleofasciculaceae, a monophyletic home for the Microcoleus steenstrupii complex and other desiccation-tolerant filamentous cyanobacteria. Journal of Phycology.

Fernandes VM, Machado de Lima NM, Roush D, Rudgers J, Collins SL & Garcia-Pichel F (2018) Exposure to predicted precipitation patterns decreases population size and alters community structure of cyanobacteria in biological soil crusts from the Chihuahuan Desert. Environmental microbiology 20: 259–269.

Field CB, Behrenfeld MJ, Randerson JT & Falkowski P (1998) Primary production of the biosphere: integrating terrestrial and oceanic components. science 281: 237–240.

Fierer N, Schimel JP & Holden PA (2003) Variations in microbial community composition through two soil depth profiles. Soil Biology and Biochemistry 35: 167–176.

Fleming ED & Castenholz RW (2007) Effects of periodic desiccation on the synthesis of the UV-screening compound, scytonemin, in cyanobacteria. Environmental Microbiology 9: 1448–1455.

Gao Q & Garcia-Pichel F (2011) Microbial ultraviolet sunscreens. Nature Reviews Microbiology 9: 791.

Garcia-Pichel F & Belnap J (2001) Small-scale environments and distribution of biological soil crusts. Biological soil crusts: Structure, function, and management,p.^pp. 193–201. Springer.

Garcia-Pichel F & Wojciechowski MF (2009) The evolution of a capacity to build supra-cellular ropes enabled filamentous cyanobacteria to colonize highly erodible substrates. PLoS One 4: e7801.

Garcia-Pichel F, Johnson S, Youngkin D & Belnap J (2003) Small-scale vertical distribution of bacterial biomass and diversity in biological soil crusts from arid lands in the Colorado Plateau. Microbial Ecology 46: 312–321.

Garcia-Pichel F, Lombard J, Soule T, Dunaj S, Wu SH & Wojciechowski MF (2019) Timing the evolutionary advent of cyanobacteria and the later great oxidation event using gene phylogenies of a sunscreen. Mbio 10: e00561–00519.

Garcia-Pichel F & Castenholz RW (1991) Characterization and biological implications of scytonemin, a cyanobacterial sheath pigment 1. Journal of Phycology 27: 395–409.

Garcia-Pichel F, Sherry ND & Castenholz RW (1992) Evidence for an ultraviolet sunscreen role of the extracellular pigment scytonemin in the terrestrial cyanobacterium Chiorogloeopsis sp. Photochemistry and photobiology 56: 17–23.

Garcia-Pichel F, Kühl M, Nübel U & Muyzer G (1999) Salinity-dependent limitation of photosynthesis and oxygen exchange in microbial mats. Journal of Phycology 35: 227–238.

Goberna M, Navarro-Cano JA, Valiente-Banuet A, García C & Verdú M (2014) Abiotic stress tolerance and competition-related traits underlie phylogenetic clustering in soil bacterial communities. Ecology letters 17: 1191–1201.

Gremer JR & Sala A (2013) It is risky out there: the costs of emergence and the benefits of prolonged dormancy. Oecologia 172: 937–947.

Hart T, Chamberlain A, Lynch J, Newling B & McDonald P (1999) A stray field magnetic resonance study of water diffusion in bacterial exopolysaccharides. Enzyme and microbial technology 24: 339–347.

Hector RF & Laniado-Laborin R (2005) Coccidioidomycosis—A Fungal Disease of the Americas. PLOS Medicine 2: e2.

Hoover DL, Bestelmeyer B, Grimm NB, Huxman TE, Reed SC, Sala O, Seastedt TR, Wilmer H & Ferrenberg S (2020) Traversing the wasteland: a framework for assessing ecological threats to drylands. BioScience 70: 35–47.

Huenneke LF & Mooney HA (2012) Grassland structure and function: California annual grassland. Springer Science & Business Media.

Jackson R (1999) The importance of root distributions for hydrology, biogeochemistry, and ecosystem functioning. Integrating Hydrology, Ecosystem Dynamics, and Biogeochemistry in Complex Landscapes 217–238.

Jakubczyk D, Kolwas M, Derkachov G, Kolwas K & Zientara M (2012) Evaporation of microdroplets: the” radius-square-law” revisited. Acta Physica Polonica-Series A General Physics 122: 709.

Jones SK, Collins SL, Blair JM, Smith MD & Knapp AK (2016) Altered rainfall patterns increase forb abundance and richness in native tallgrass prairie. Scientific reports 6: 1–10.

Jose NA, Lau R, Swenson TL, Klitgord N, Garcia-Pichel F, Bowen BP, Baran R & Northen TR (2018) Flux balance modeling to predict bacterial survival during pulsed-activity events. Biogeosciences 15: 2219–2229.

Karaoz U, Couradeau E, Da Rocha UN, Lim H-C, Northen T, Garcia-Pichel F & Brodie EL (2018) Large blooms of Bacillales (Firmicutes) underlie the response to wetting of cyanobacterial biocrusts at various stages of maturity. mBio 9: e01366–01316.

Klotz A & Forchhammer K (2017) Glycogen, a major player for bacterial survival and awakening from dormancy. p.^pp. Future Medicine.

Klotz A, Georg J, Bučinská L, Watanabe S, Reimann V, Januszewski W, Sobotka R, Jendrossek D, Hess WR & Forchhammer K (2016) Awakening of a dormant cyanobacterium from nitrogen chlorosis reveals a genetically determined program. Current Biology 26: 2862–2872.

Kolodny NH, Bauer D, Bryce K, Klucevsek K, Lane A, Medeiros L, Mercer W, Moin S, Park D & Petersen J (2006) Effect of nitrogen source on cyanophycin synthesis in Synechocystis sp. strain PCC 6308. Journal of bacteriology 188: 934–940.

Lauenroth W, Sala O, Milchunas D & Lathrop R (1987) Root dynamics of Bouteloua gracilis during short-term recovery from drought. Functional Ecology 117–124.

Le PT, Makhalanyane TP, Guerrero LD, Vikram S, Van de Peer Y & Cowan DA (2016) Comparative Metagenomic Analysis Reveals Mechanisms for Stress Response in Hypoliths from Extreme Hyperarid Deserts. Genome biology and evolution 8: 2737–2747.

LeBlanc JC, Gonçalves ER & Mohn WW (2008) Global response to desiccation stress in the soil actinomycete Rhodococcus jostii RHA1. Applied and environmental microbiology 74: 2627–2636.

Lennon JT & Jones SE (2011) Microbial seed banks: the ecological and evolutionary implications of dormancy. Nature reviews microbiology 9: 119–130.

Lennon JT, den Hollander F, Wilke-Berenguer M & Blath J (2021) Principles of seed banks and the emergence of complexity from dormancy. Nature Communications 12: 1–16.

Linn DM & Doran JW (1984) Effect of water-filled pore space on carbon dioxide and nitrous oxide production in tilled and nontilled soils 1. Soil Science Society of America Journal 48: 1267–1272.

Liu D, Keiblinger KM, Leitner S, Wegner U, Zimmermann M, Fuchs S, Lassek C, Riedel K & Zechmeister-Boltenstern S (2019) Response of Microbial Communities and Their Metabolic Functions to Drying–Rewetting Stress in a Temperate Forest Soil. Microorganisms 7: 129.

Madueño L, Coppotelli BM, Festa S, Alvarez H & Morelli IS (2018) Insights into the mechanisms of desiccation resistance of the Patagonian PAH-degrading strain Sphingobium sp. 22B. Journal of applied microbiology 124: 1532–1543.

Martínez-Vilalta J, Sala A, Asensio D, Galiano L, Hoch G, Palacio S, Piper FI & Lloret F (2016) Dynamics of non-structural carbohydrates in terrestrial plants: a global synthesis. Ecological Monographs 86: 495–516.

Martiny JB, Jones SE, Lennon JT & Martiny AC (2015) Microbiomes in light of traits: a phylogenetic perspective. Science 350.

Mason-Jones K, Robinson SL, Veen G, Manzoni S & van der Putten WH (2021) Microbial storage and its implications for soil ecology. The ISME Journal 1–13.

Menke JW & Trlica M (1983) Effects of single and sequential defoliations on the carbohydrate reserves of four range species. Rangeland Ecology & Management/Journal of Range Management Archives 36: 70–74.

Neilson JW, Califf K, Cardona C, et al. (2017) Significant Impacts of Increasing Aridity on the Arid Soil Microbiome. mSystems 2: e00195–00116.

Noy-Meir I (1973) Desert ecosystems: environment and producers. Annual review of ecology and systematics 4: 25–51.

Ogle K & Reynolds JF (2004) Plant responses to precipitation in desert ecosystems: integrating functional types, pulses, thresholds, and delays. Oecologia 141: 282–294.

Oren N, Raanan H, Kedem I, Turjeman A, Bronstein M, Kaplan A & Murik O (2019) Desert cyanobacteria prepare in advance for dehydration and rewetting: The role of light and temperature sensing. Molecular ecology 28: 2305–2320.

Orians GH & Solbrig OT (1977) A cost-income model of leaves and roots with special reference to arid and semiarid areas. The American Naturalist 111: 677–690.

Orians GH & Solbrig OT (1977) Convergent evolution in warm deserts. An examination of strategies and patterns in deserts of Argentina and the United States. US/IBP Synthesis Series (USA) v 3.

Planinšič G & Vollmer M (2008) The surface-to-volume ratio in thermal physics: from cheese cube physics to animal metabolism. European Journal of Physics 29: 369.

Potts M (2001) Desiccation tolerance: a simple process? Trends in microbiology 9: 553–559.

Poulter B, Frank D, Ciais P, Myneni RB, Andela N, Bi J, Broquet G, Canadell JG, Chevallier F & Liu YY (2014) Contribution of semi-arid ecosystems to interannual variability of the global carbon cycle. Nature 509: 600–603.

Prăvălie R (2016) Drylands extent and environmental issues. A global approach. Earth-Science Reviews 161: 259–278.

Price C & Munns R (2020) Chapter 6—Growth analysis: A quantitative approach. Plants in action 24.

Pringault O & Garcia-Pichel F (2004) Hydrotaxis of cyanobacteria in desert crusts. Microbial ecology 47: 366–373.

Rajeev L, da Rocha UN, Klitgord N, et al. (2013) Dynamic cyanobacterial response to hydration and dehydration in a desert biological soil crust. The Isme Journal 7: 2178.

Reed SC, Coe KK, Sparks JP, Housman DC, Zelikova TJ & Belnap J (2012) Changes to dryland rainfall result in rapid moss mortality and altered soil fertility. Nature Climate Change 2: 752–755.

Reynolds JF, Smith DMS, Lambin EF, Turner B, Mortimore M, Batterbury SP, Downing TE, Dowlatabadi H, Fernández RJ & Herrick JE (2007) Global desertification: building a science for dryland development. science 316: 847–851.

Richardson DM, Hellmann JJ, McLachlan JS, Sax DF, Schwartz MW, Gonzalez P, Brennan EJ, Camacho A, Root TL & Sala OE (2009) Multidimensional evaluation of managed relocation. Proceedings of the National Academy of Sciences 106: 9721–9724.

Roberson EB & Firestone MK (1992) Relationship between desiccation and exopolysaccharide production in a soil Pseudomonas sp. Appl Environ Microbiol 58: 1284–1291.

Rothrock Jr MJ & Garcia-Pichel F (2005) Microbial diversity of benthic mats along a tidal desiccation gradient. Environmental Microbiology 7: 593–601.

Rothschild LJ & Mancinelli RL (2001) Life in extreme environments. Nature 409: 1092.

Sala O, Lauenroth W & Parton W (1992) Long-term soil water dynamics in the shortgrass steppe. Ecology 73: 1175–1181.

Scherer S & Zhong Z-P (1991) Desiccation independence of terrestrialNostoc commune ecotypes (Cyanobacteria). Microbial ecology 22: 271–283.

Schimel JP (2018) Life in dry soils: effects of drought on soil microbial communities and processes. Annual review of ecology, evolution, and systematics 49: 409–432.

Schlesinger WH, Pippen JS, Wallenstein MD, Hofmockel KS, Klepeis DM & Mahall BE (2003) Community composition and photosynthesis by photoautotrophs under quartz pebbles, southern Mojave Desert. Ecology 84: 3222–3231.

Schulz H, Brinkhoff T, Ferdelman T, Mariné MH, Teske A & Jørgensen B (1999) Dense populations of a giant sulfur bacterium in Namibian shelf sediments. Science 284: 493–495.

Schulze-Makuch D, Wagner D, Kounaves SP, Mangelsdorf K, Devine KG, de Vera J-P, Schmitt-Kopplin P, Grossart H-P, Parro V & Kaupenjohann M (2018) Transitory microbial habitat in the hyperarid Atacama Desert. Proceedings of the National Academy of Sciences 115: 2670–2675.

Schwinning S, Sala OE, Loik ME & Ehleringer JR (2004) Thresholds, memory, and seasonality: understanding pulse dynamics in arid/semi-arid ecosystems. p.^pp. Springer.

Setlow P (2007) I will survive: DNA protection in bacterial spores. Trends in microbiology 15: 172–180.

Simpson EL, Heness E, Bumby A, Eriksson PG, Eriksson KA, Hilbert-Wolf HL, Linnevelt S, Malenda HF, Modungwa T & Okafor O (2013) Evidence for 2.0 Ga continental microbial mats in a paleodesert setting. Precambrian Research 237: 36–50.

Smith S, Abed RMM & Gercia-Pichel F (2004) Biological soil crusts of sand dunes in Cape Cod National Seashore, Massachusetts, USA. Microbial ecology 48: 200–208.

Stewart G, Johnstone K, Hagelberg E & Ellar DJ (1981) Commitment of bacterial spores to germinate A measure of the trigger reaction. Biochemical Journal 198: 101–106.

Šťovíček A, Kim M, Or D & Gillor O (2017) Microbial community response to hydration-desiccation cycles in desert soil. Scientific reports 7: 1–9.

Sukenik A, Kaplan-Levy RN, Welch JM & Post AF (2012) Massive multiplication of genome and ribosomes in dormant cells (akinetes) of Aphanizomenon ovalisporum (Cyanobacteria). The ISME Journal 6: 670–679.

Sukenik A, Maldener I, Delhaye T, Viner-Mozzini Y, Sela D & Bormans M (2015) Carbon assimilation and accumulation of cyanophycin during the development of dormant cells (akinetes) in the cyanobacterium Aphanizomenon ovalisporum. Frontiers in microbiology 6: 1067.

Thomazo C, Couradeau E & Garcia-Pichel F (2018) Possible nitrogen fertilization of the early Earth Ocean by microbial continental ecosystems. Nature Communications 9: 2530.

Warren-Rhodes KA, Rhodes KL, Pointing SB, Ewing SA, Lacap DC, Gomez-Silva B, Amundson R, Friedmann EI & McKay CP (2006) Hypolithic cyanobacteria, dry limit of photosynthesis, and microbial ecology in the hyperarid Atacama Desert. Microbial ecology 52: 389–398.

Weissman JL, Hou S & Fuhrman JA (2021) Estimating maximal microbial growth rates from cultures, metagenomes, and single cells via codon usage patterns. Proceedings of the National Academy of Sciences 118.

Yahdjian L, Sala OE & Havstad KM (2015) Rangeland ecosystem services: shifting focus from supply to reconciling supply and demand. Frontiers in Ecology and the Environment 13: 44–51.

